# Targeting Tuberculosis: Novel Scaffolds for Inhibiting Cytochrome bd Oxidase

**DOI:** 10.1101/2024.02.28.582612

**Authors:** Christian Seitz, Surl-Hee Ahn, Haixin Wei, Matson Kyte, Gregory M. Cook, Kurt Krause, J. Andrew McCammon

**Author notes:** These authors have contributed equally (Christian Seitz, Surl-Hee Ahn, Haixin Wei, Matson Kyte).

## Abstract

Discovered in the 1920s, cytochrome *bd* is a terminal oxidase that has received renewed attention as a drug target since its atomic structure was first solved in 2016. Only found in prokaryotes, we study it here as a drug target for *Mycobacterium tuberculosis* (*Mtb*). Most previous drug discovery efforts towards cytochrome *bd* have involved analogs of the canonical substrate quinone, known as Aurachin D. Here we report six new cytochrome *bd* inhibitor scaffolds determined from a computational screen totaling over one million molecules and confirmed on target activity through *in vitro* testing. These scaffolds provide new avenues for lead optimization towards *Mtb* therapeutics.

## Introduction

Cytochrome *bd* oxidase, or cytochrome *bd*,^1^ is an oxygen reduction enzyme found only in prokaryotes that reduces oxygen to water in the aerobic respiration cycle. Ubiquinol (or menaquinol) binds to cytochrome *bd* and is oxidized to ubiquinone (or menaquinone).^2^ Combined with proton binding, this allows molecular oxygen to be reduced to water.^2^ Cytochrome *bd* contains three hemes; the roles and organization of these hemes differ across organisms.^3^ The few known cytochrome *bd* structures have contained nine to twenty transmembrane helices, depending on the organism. ^1-3^

The location of cytochrome *bd* in the bacterial cytoplasmic membrane allows it to bind ubiquinol, or menaquinol in the case of *Mycobacterium tuberculosis* (*Mtb*), and oxygen in the periplasm and accept free protons in the cytoplasm.^2^ It contains two to four subunits, depending on the organism.^1^ It was first described in 1928,^4^ but no structure was solved from any bacteria until 2016 when one structure was solved in *Geobacillus thermodenitrificans*.^5^ Since then, four structures have been solved from *Escherichia coli (E. coli)*,^6-9^ one from *Mtb*,^10^ and one from *Mycobacterium smegmatis*.^11^

Cytochrome *bd* is an attractive drug target for two reasons. The first is that it appears to be an important feature of many pathogenic prokaryotes; for example, it is found in Mycobacterium tuberculosis, which causes tuberculosis; Vibrio cholerae, which causes cholera; Pseudomonas aeruginosa, which contributes to antibiotic resistance and sepsis; *Campylobacter jejuni*, which is a common cause of food poisoning, among many other disease-causing bacteria. The second is that cytochrome *bd* is not found in humans, which would reduce the chance of drug-inducing side effects. Drug discovery efforts towards cytochrome *bd* have been aided by the elucidation of cytochrome *bd* structures. Considering that quinones are cytochrome *bd’s* biological substrate, it is unsurprising that most of the potential inhibitors so far have come from quinone analogs.^6-10, 12-19^ One group screened a diverse molecular library for cytochrome *bd* inhibitors, with the resulting hits being more quinone analogs^20^ whereas only a few other scaffolds have been identified as inhibitors.^21-24^ These inhibitors have been reviewed elsewhere. ^25^

Although cytochrome *bd* inhibitors are at a very early stage of drug development, they are an attractive addition to a future multidrug cocktail that could be used to treat tuberculosis, including drug-resistant isolates.^26^ To further develop new cytochrome *bd* inhibitor scaffolds, we ran molecular dynamics (MD) simulations of cytochrome *bd* in both Mtb and *E. coli* to extract a structural variety of potential inhibitor pockets or surface-level binding sites. Then, we screened ∼1 million molecules in these regions through molecular docking before testing the most promising compounds against *Mtb* cytochrome bd *in vitro*.

## Methods

### Gaussian accelerated molecular dynamics & simulation setup

Gaussian accelerated molecular dynamics (GaMD) is an enhanced sampling method for MD simulations that can efficiently sample various conformations of the protein of interest.^27-29^ GaMD gains its efficiency by filling the energy wells with a harmonic boost potential Δ*V* (**r**), where **r** denotes the position vector of an *N*-atom system and *V* (**r**) denotes the system potential, i.e.,

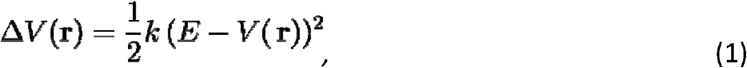

where *k* denotes the harmonic force constant, and E denotes the reference energy. Δ*V* (**r**) is only added if *V* (**r**) < *E*. Specific details on how Δ*V* (**r**), *E*, and *k* are set are stated in the original GaMD paper.^27-29^ GaMD was primarily used here to generate conformational ensembles for virtual screening and docking.

Three initial cytochrome *bd* structures for the GaMD simulations were prepared: the *Escherichia coli (E. coli)* cytochrome *bd* (PDB: 6RKO), the *Mtb* cytochrome *bd* (PDB: 7NKZ) with the disulfide bond within the quinol-binding and oxidation domain (Q-loop) intact, which defines an inactive conformation, and the *Mtb* cytochrome *bd* (PDB: 7NKZ) with the disulfide bond broken, which defines an active conformation.^6, 10^ *E. coli* cytochrome *bd* has the highest sequence identity to that of *Mtb* (36 %), which makes it a suitable system for virtual screening and docking.^6, 30, 31^ Schrödinger Maestro was used to add hydrogens to the residues and assign protonation states to the residues using PROPKA at pH 7.4.^32^ The AlphaFold structures from UniProt were used for *E. coli* cytochrome *bd* and *Mtb* cytochrome *bd* systems.^33^ The Amber 20 MD simulation engine was used to run the MD simulations for each system.^34^

The lipid bilayer for the *E. coli* cytochrome *bd* oxidase system was constructed through CHARMM-GUI^35, 36^ and was based on the experimentally determined composition.^37^ Our setup contained 79% phosphatidylethanolamine (PE) lipids (5% 1,2-dimyristoyl-sn-glycero-3-phosphoethanolamine [DMPE], 40% 1,2-dipalmitoyl-sn-glycero-3-phosphoethanolamine [DPPE], 17% 1-palmitoyl-2-oleoylphosphatidylethanolamine [POPE], 17% 1,2-dioleoyl-sn glycero-3phosphoethanolamine [DOPE]) and 21% phosphatidylglycerol (PG) lipids (1% 1,2 dimyristoylsn-glycero-3-phosphoglycerol, 14% 1,2-dipalmitoyl-sn-glycero-3 phosphoglycerol, 6% 1,2-dioleoylsn-glycero-3-phosphoglycerol [DOPG]). The lipid bilayer for the *Mtb* cytochrome *bd* oxidase system was also constructed through CHARMM-GUI^35, 36^ and was based on the experimentally determined composition.^38-41^ The setup for this bilayer contained 98% PE lipids (5% DMPE, 31% DPPE, 31% POPE, 31% DOPE) and 2% PG lipids (2% DOPG).

Besides the lipid compositions, the systems were prepared identically except for the ion concentration noted. After insertion of the protein, each system was solvated with TIP3P.^42^ The *E. coli* cytochrome *bd* system was neutralized and had an ionic concentration of 0.10 M NaCl, while the *Mtb* cytochrome *bd* system was neutralized and had an ionic concentration of 0.15 M NaCl.^43^ Input files for the Amber MD simulations were generated using the Amber ff14SB,^44^ Lipid 17,^45^ TIP3P, ^42^ and GAFF parameters.^46^ This was done on each system without the hemes since the heme parameters were unavailable on CHARMM-GUI. Thus, after the lipid bilayer system was created, the exact same protein with the hemes was aligned to the C_α_ atoms of the heme-less protein situated in the lipid bilayer in PyMOL.^47^ This generated the correct spatial coordinates, and the heme-less protein situated in the lipid bilayer was deleted before the protein containing the hemes and the lipid bilayer were concatenated to create the systems used going forward. The hemes used our previously developed custom force field parameters.^30^

The equilibrated structures for each system were obtained by first minimizing the system for 10,000 steps with the backbone atoms and heavy atoms of the heme cofactors restrained with a force constant of 10 kcal/(mol × Å^2^). Then, each system was minimized for 100,000 steps with no restraints and gradually heated from 10 to 300 K over 4 ns and kept at 300 K and 1 bar for 6 ns in the NPT ensemble (using a Langevin thermostat with a friction coefficient *γ* of 5.0 ps^−1^ and a Berendsen barostat with a time constant *τ* of 1.0 ps) with the backbone atoms and heavy atoms of the heme cofactors restrained with a force constant of 10.0 kcal/(mol × Å^2^) for the gradual heating stage and 1.0 kcal/(mol × Å^2^) for the latter heating stage. Finally, each system was equilibrated for 10 ns in the NPT ensemble at 300 K and 1 bar (using a Langevin thermostat with a friction coefficient *γ* of 1.0 ps^−1^ and a Berendsen barostat with a time constant *τ* of 1.0 ps) with the backbone atoms and heavy atoms of the heme cofactors restrained with a force constant of 0.1 kcal/(mol × Å^2^). After equilibration, a 100 ns conventional production MD simulation was run in the NPT ensemble at 300 K and 1 bar (using a Langevin thermostat with a friction coefficient *γ* of 1.0 ps^−1^ and a Berendsen barostat with a time constant *τ* of 1.0 ps) with no restraints for each system. The root-mean-square deviation (RMSD) of all C_α_ atoms from the starting structure for the conventional production MD run plateaued during the 100 ns conventional production MD run for all systems (Figures S1).

Using the last frame from the 100 ns conventional production MD simulation run as the starting structure, six GaMD simulations were run for 500 ns each with different boost potentials (i.e., boost on total energy with the threshold energy set to upper (E = V_min_ + (V_max_ – V_min_)/k_0_, where V_max_ and V_min_ are the minimum and maximum potential energies of the system, and k_0_ is the magnitude of the applied boost potential) or lower bound (E = V_max_), boost on dihedral energy with the threshold energy set to upper/lower bound, and boost on both total and dihedral energies with the threshold energy set to upper/lower bound) for each system. For GaMD simulations, the lipid bilayer was stripped to save computational cost, and two minimization runs were done before the simulations were run in the NVT ensemble at 300 K (using a Langevin thermostat with a friction coefficient *γ* of 1.0 ps^−1^). First, the system was minimized for 10,000 steps with the backbone and heavy atoms of the heme cofactors restrained with a force constant of 10 kcal/(mol × Å^2^). Then, the system was minimized for 100,000 steps with no restraints. With one 100 ns conventional production MD simulation and six 500 ns GaMD simulations and using every 0.5 ns simulation frame of the combined 600 ns trajectory for each system, clustering was performed for each binding site to get representative conformations for each binding site. The binding sites were reported in the literature as potential binding sites to target for drug development (Figure S1).^5,10^ For the *E. coli* cytochrome *bd*, the entry site of the oxygen conducting channel (which starts from Trp 63.B), the Q-loop (Lys 250.A to Arg 382.A), and the ubiquinol binding site (between transmembrane helix 6 and transmembrane helix 7 in CydB subunit as noted in Figure 1 of Ref. 6) were the possible binding sites. For the *Mtb* cytochrome *bd* with and without the disulfide bond, the entry site of the oxygen conducting channel (which starts from Trp 63.B), the Q-loop (Pro 256.A to Asn 333.A), menaquinone binding site (which consists of the heme b595 porphyrin scaffold, Arg8.A and Trp9.A of transmembrane helix 1 in CydA subunit, and Met397.A of transmembrane helix 9 in CydA subunit), and the region between the C-terminal region of the Q-loop (QC) and the large periplasmic loop connecting transmembrane helices 8 and 9 (PL8) (Thr 310.A to Asn 333.A and Gln 400.A to Ala 420.A) were the possible binding sites. The DBScan clustering algorithm from the Amber CPPTRAJ program was used to cluster the Q-loop and the region between QC and PL8. In contrast, the POcket Volume MEasurer (POVME) 2.0 was used to cluster the other sites since the other sites were not in the outermost, solvent-exposed regions, unlike the Q-loop and the region between QC and PL8.^48, 49^ The POVME calculations were started from the center of the pocket defined according to the aforementioned binding site definitions. Using POVME, the volumes of each binding site or pocket were calculated from the combined 600 ns trajectory, and the simulation frames that were representative of the “peaks” or points where a particular volume was sampled most frequently were extracted to be one of the representative conformations (Figures 1 and 2). These representative conformations were then used in the molecular docking section.

**Figure 1.**
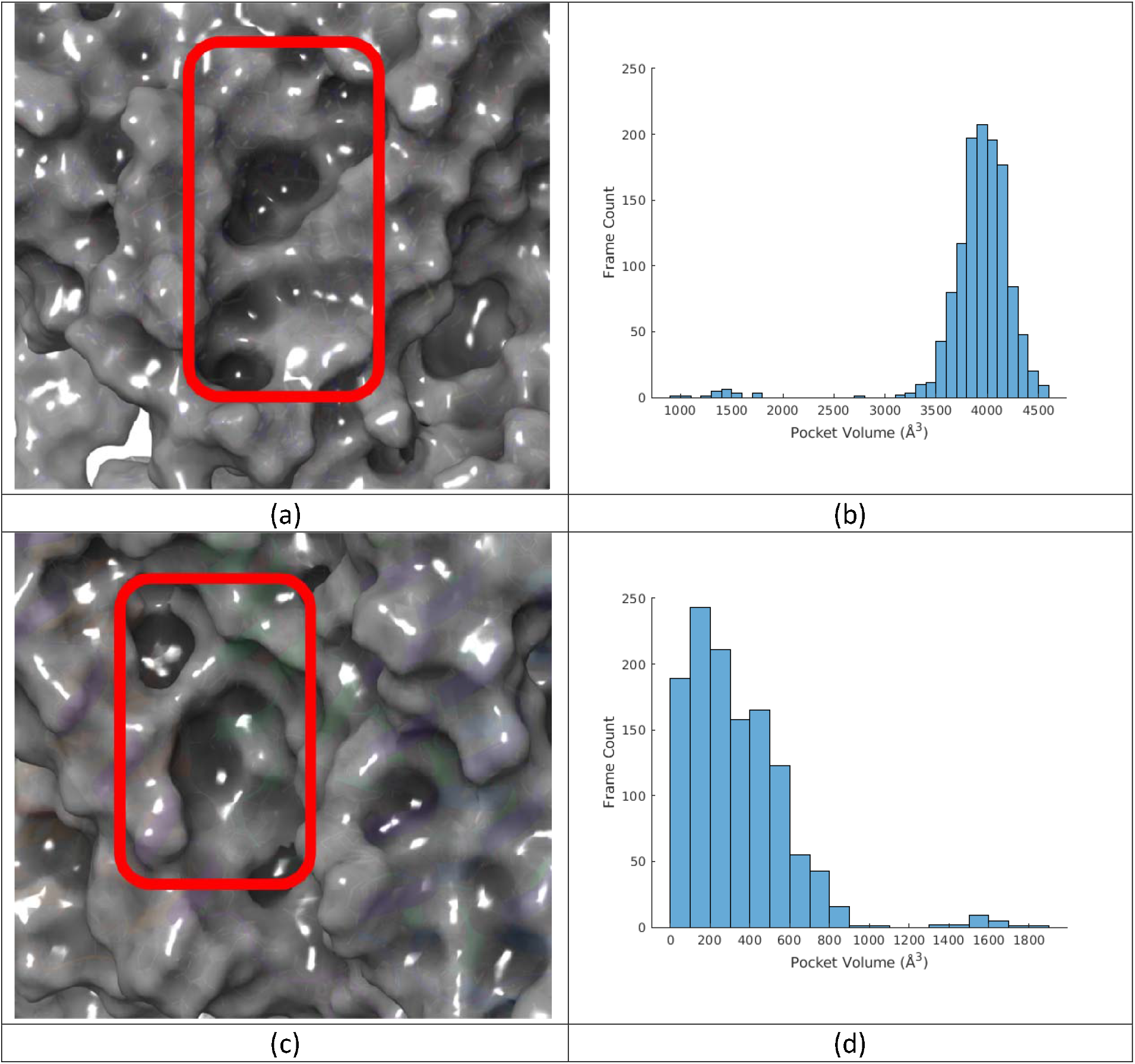
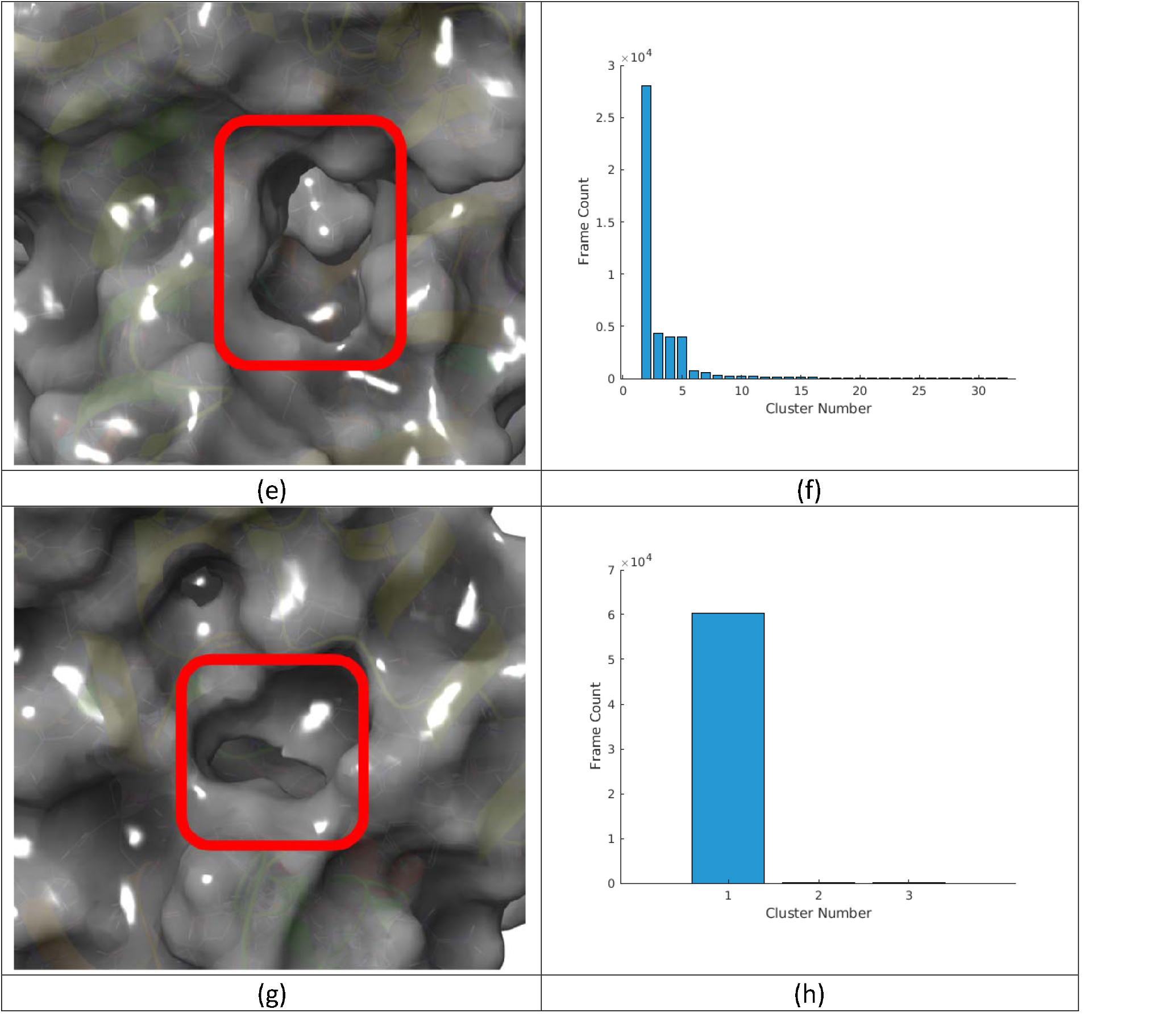
Representative pocket conformations and the population distribution for GaMD trajectories of *Mtb* cytochrome *bd* without the disulfide bond (active conformation). (a) MQ9 binding site and (b) pocket volume distribution of it; (c) oxygen conducting channel and (d) pocket volume distribution of it; (e) the Q-loop region and (f) cluster population distribution of it; (g) the region between QC and PL8 and (h) cluster population distribution of it.

**Figure 2.**
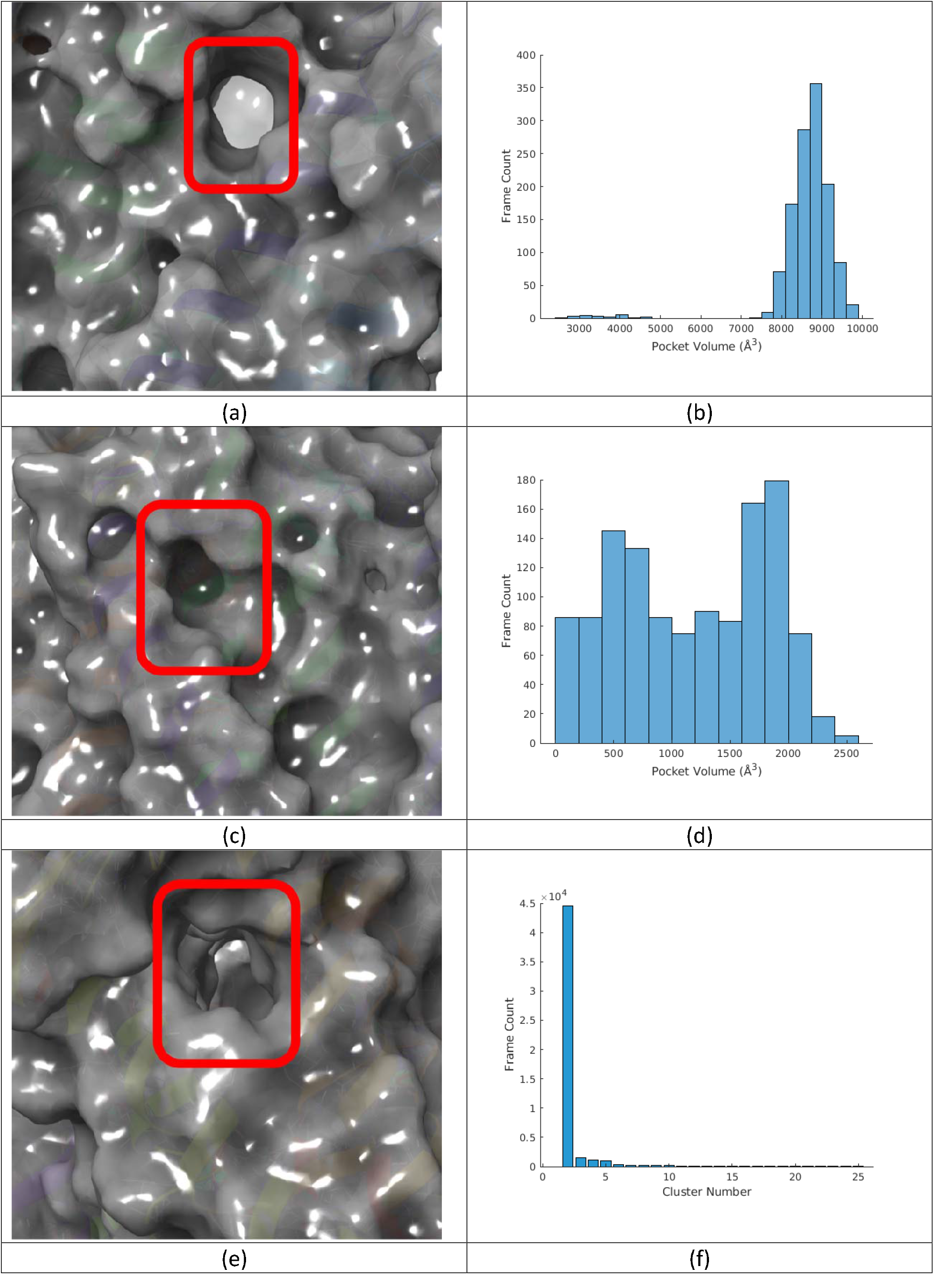
Representative pocket conformations and the population distribution for GaMD trajectory of *E. coli* cytochrome *bd*. (a) UQ8 binding site and (b) pocket volume distribution of it; (c) oxygen conducting channel and (d) pocket volume distribution of it; (e) the Q-loop region and (f) cluster population distribution of it.

For the *E. coli* cytochrome *bd* system, the following conformations were selected and used in the molecular docking section. The four most representative clusters in terms of the number of simulation frames were used for the Q-loop. Conformations with pocket volumes of 598 Å, 1170 Å, and 1846 Å were used for the oxygen conducting channel, and conformations with pocket volumes of 3538 Å and 8646 Å were used for the ubiquinol binding site (Figure 2). For the *Mtb* cytochrome *bd* with the disulfide bond, the following conformations were selected and used in the molecular docking section. The four most representative clusters in terms of the number of simulation frames were used for the Q-loop, and the top representative cluster was used for the region between QC and PL8. Conformations with pocket volumes of 175 °Å^3^, 351 °Å^3^, 825 Å^3^, and 1845 Å^3^ were used for the oxygen conducting channel, and conformations with pocket volumes of 4000 Å^3^ and 8244 Å^3^ were used for the menaquinone binding site. For the *Mtb* cytochrome *bd* without the disulfide bond, the following conformations were selected and used in the molecular docking section. The four most representative clusters in terms of the number of simulation frames were used for the Q-loop, and the three most representative clusters were used for the region between QC and PL8. Conformations with pocket volumes of 190 Å^3^, 336 Å^3^, 437 Å^3^, and 1535 Å^3^ were used for the oxygen conducting channel, and conformations with pocket volumes of 1412 Å^3^ and 3919 Å^3^ were used for the menaquinone binding site (Figure 1).

### Molecular docking

Each structure generated from the clustering stage was put through the same general docking workflow, with differences noted below. First, the structure was run through Schrödinger’s Protein Preparation Wizard.^48^ This involved capping the termini, filling in missing side chains, assigning bond orders using the Cambridge crystallographic database, creating zero-order bonds to metals, creating disulfide bonds, filling in missing loops using Prime, and generating het states with Epik using a pH of 7.4 ± 2.0.^50, 51^ We did not optimize hydrogen bond assignments or clean up the structure, as these steps would have altered the structures generated from the clustered GaMD work.

The binding site definitions for each site are the same as those described previously unless otherwise noted. The receptor grids were created in Schrödinger Maestro.^31^ For the *Mtb* Q-loop with the disulfide bond, we moved the bounding box 10 Å in the -Y direction to better fit over the open space near the Q-loop. This was done for each of the four protein structures used. For the *Mtb* Q-loop without the disulfide bond, we only moved the bounding box 15 Å in the +X direction for the structure from the first cluster and 5 Å in the +X direction for the structures from the second and fourth clusters. For *E. coli*, we docked ligands with length ≤ 18 Å, while for Mtb, we docked ligands with length ≤ 14 Å; these values were chosen so that docked poses would fit within their respective protein structures without undue steric clashes. The box length in X and Y was 40 Å, while the box length in Z was 28 Å for both systems. The UQ8 region in *E. coli* and the MQ9 region in *Mtb* region were defined as being a box around the crystallized, bound ligand, docking ligands the same size as either UQ8 or MQ9. We increased the *Mtb* oxygen channel box coordinates to +15 Å in each direction.

The first library we used for docking was the Enamine Advanced Collection, which contained 640,219 compounds when downloaded on March 17, 2022. This collection is curated to contain compounds synthesized within the last three years with structural trends in medicinal chemistry designs. The second library was the Life Chemicals HTS Compound Collection, which contained 523,192 compounds when downloaded on March 21, 2022. This library contains structurally diverse compounds with optimized physicochemical properties for drug discovery projects. When needed, the solvent, crystallized ligands, and hemes were deleted from the input structures as these did not participate in docking. The Enamine Advanced Collection and the Life Chemicals HTS Compound Collection were separately docked to each protein structure selected from the clustering. These compounds were then docked using Schrödinger Glide’s HTVS functionality,^52^ which does a rough structure-based energy screening of the ligand fitting into the protein’s binding pocket. Then, the top 20,000 scoring poses were carried forward to the next selection round. Note that these were 20,000 poses, not necessarily 20,000 unique ligands, so some ligands were represented in several poses that were all carried forward. Next, these top 20,000 scoring poses were docked using Schrödinger Glide’s SP module, which uses the same scoring function as HTVS but increases the number of intermediate conformations throughout the docking funnel and increases torsional refinement and sampling. Finally, the top 1,000 poses were carried forward to Schrödinger Glide’s XP docking, which uses a more sophisticated scoring function than HTVS or SP, reducing the number of false positives. At this point, the top *n*-scoring number of ligands, not the top *n*-number of poses, were selected for experimental screening. For the *E. coli* and *Mtb* work, 50 molecules from Enamine and Life Chemicals were purchased for testing, totaling 200 molecules. Thus, this workflow meant that all molecules could be cross-docked to all possible structures, with the top-scoring structures tested experimentally.

### Experimental Testing

In preparation of inverted membrane vesicles expressing the *M. tuberculosis* cytochrome *bd*, strain *M. smegmatis* Δ*cyd* pLHcyd was used as previously described^10^. All cultures were grown in LB media (10 g/L tryptone, 5 g/L yeast extract, 5 g/L NaCl) with 100 µg/mL of hygromycin B added to media before use, with agitation at 200 RPM and maintained at 37 °C.

To produce IMVs for biochemical assays, a single colony of *M. smegmatis* Δ*cyd* pLHcyd was selected to inoculate a 5 mL starter culture. After 72 h of growth, this culture was used to inoculate 1 L of production culture grown in a 2 L flask in a 1:200 dilution. The production culture was grown for 72 h before cells were harvested by centrifugation at 4000 × g for 20 min at 4 °C. Harvested cells were frozen at -80 °C. Cells were resuspended and thawed at a ratio of 1 g of wet-weight cells to 5 mL of buffer containing 50 mM Tris-HCl (pH 7.4), 5 mM MgCl2, and 1 cOmplete™ mini protease inhibitor cocktail tablet per 50 mL. Once fully thawed and resuspended, cells were passed four to six times through a high-pressure homogenizer at 22,000 psi with a 5-minute rest on ice between passes. The resulting cell lysate was centrifuged at 15,000 × g for 10 mins at 4 °C to remove cell debris and unlysed cells. The supernatant from this spin was decanted and centrifuged again at 200,000 × g for 45 min at 4 °C to separate the cytosolic and membrane fractions. The pelleted membranes were resuspended in a buffer containing 50⍰mM Tris-HCL (pH 7.4), 100⍰mM KCl, 5⍰mM MgCl2, and 10% glycerol. Before freezing at -80 °C, the protein concentration was estimated by BCA assay (Thermo).

OCR was measured using an Oroboros O2k FluoRespirometer as previously described^24^. Assay buffer (2 mL) containing 50 mM Tris-HCL (pH 7.0), 5 mM MgCl_2_, and 100 mM KCl was added to measurement chambers. IMVs were added at a concentration of 75 µg/mL along with 1 µM TB47 for approximately 2 min to inhibit the activity of the *bcc:aa*_3_ oxidase complex^53^ NADH (500 µM) was added to initiate respiration. Once OCR had stabilized, OCR was recorded, and a test compound/inhibitor was added at a final concentration of 10 µM. Once OCR had again stabilized, the new OCR was recorded. All compounds were tested once, and compounds that decreased OCR to below 50% of the initial measurement were selected for further testing. DMSO and Aurachin D were used as vehicle control and positive control, respectively^24^.

## Results and Discussion

### 1. Characterization of the target pockets

As previously mentioned, the binding sites were reported in the literature as potential binding sites to target for drug development (Figure S3).^5,10^ Here, in Figure 1 and Figure 2, we showed the structures of the four chosen pockets in *Mtb* cytochrome *bd* in active conformation and the three chosen pockets in *E. coli* cytochrome *bd*, respectively. Similar plot for *Mtb* cytochrome *bd* in active conformation are shown in Figure S2. We chose to investigate cytochrome *bd* in both *Mtb* and *E. coli* orthologs to diversify the drug pockets we investigated with the hopes of identifying cytochrome *bd* inhibitors across different bacterial genera that harbour cytochrome *bd*.

To derive representative conformation from our GaMD trajectories, we used two clustering methods: POVME 2.0 and the DBScan clustering algorithm from the Amber CPPTRAJ program. POVME can calculate and cluster binding pocket volumes. However, the Q-loop and QC-PL8 binding regions are binding surfaces, not defined pockets, which necessitated using the RMSD-based DBScan to cluster these solvent-exposed surfaces. Here, we use *Mtb* cytochrome *bd* system in active conformation as an example, but similar analysis can be done for other two systems. After calculating the pocket volumes of the traditional binding pockets, one can examine a histogram of those volumes and extract protein structures displaying the most representative pocket volumes. We selected pocket volumes of 3919 Å^3^ (representing the major peak seen in Figure 1(b)) and 1412 Å^3^ (representing the minor peak seen in Figure 1(b)). Similarly, we chose pocket volumes of 190 Å^3^, 336 Å^3^, and 437 Å^3^ to sample better the major peak for the oxygen conducting channel, this time neglecting the minor peak around 1500 Å^3^ (Figure 1(d)).

Figures 1(f) and 1(h) show the clustering results of the Q-loop and QC-PL8 regions. There are four representative conformations for the Q-loop region (Figure 1(f)). All other conformations are considerably less occupied and are assumed to be negligible. For the QC-PL8 pockets, the algorithm returns only three clusters. As a result, we included all of them in our ensemble.

### 2. Docking results

After choosing binding pockets or regions and clustering the MD trajectories, we eventually picked 33 structures (13 from *Mtb* cytochrome *bd* without the disulfide bond, 11 from *Mtb* cytochrome *bd* with the disulfide bond, and 9 from *E. coli* cytochrome *bd* oxidase) as our receptor ensemble for the docking process. The docking was done using the Schrödinger Maestro package, and the results with the best scores of each structure are shown in Tables 1 and 2.

**Table 1.** The docked compounds with the best scores for each of the selected conformations and pockets of *Mtb* cytochrome *bd* without disulfide bond (active conformation). (Oxy_190 stands for the pocket at the oxygen conducting channel with a volume of 190 A3; MQ9_1412 stands for the pocket at the MQ9 binding site with a volume of 1412 A3. The rest are of similar notation.) The compound names are the Life Chemicals ID numbers.

|  | Compounds | Scores | Cluster |
| --- | --- | --- | --- |
| Q-loop | F3025-0033 | -8.943 | Most occupied |
|  | F1885-0637 | -10.260 | 2<sup>nd</sup> occupied |
|  | F6782-3644 | -10.346 | 3<sup>rd</sup> occupied |
|  | F0192-0554 | -11.572 | 4<sup>th</sup> occupied |
| QC-PL8 | F0913-5493 | -12.190 | Most occupied |
|  | F6517-0436 | -11.681 | 2<sup>nd</sup> occupied |
|  | F2758-0216 | -11.111 | 3<sup>rd</sup> occupied |
| Oxygen channel | F3386-2591 | -10.675 | Oxy_190 |
|  | F1601-0481 | -11.248 | Oxy_336 |
|  | F5027-0632 | -8.772 | Oxy_437 |
|  | F0916-5053 | -9.905 | Oxy_1535 |
| MQ9 | F6548-3745 | -10.121 | MQ9_1412 |
|  | F6548-3808 | -8.220 | MQ9_3919 |

**Table 2.**
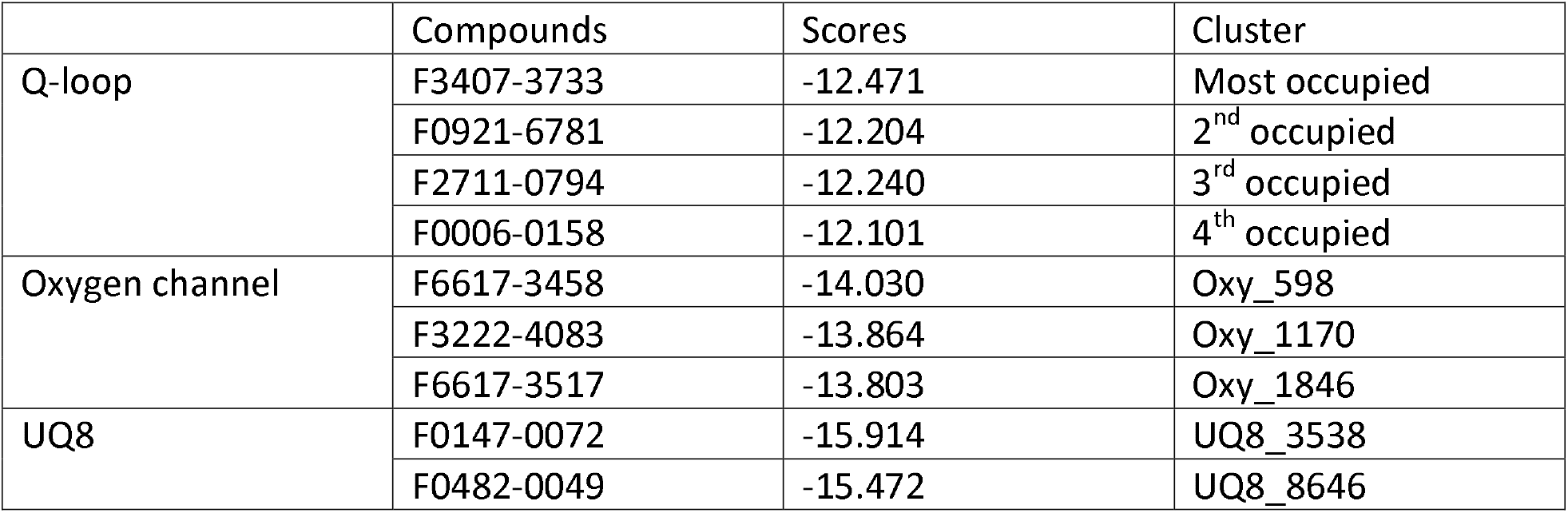
The docked compounds with the best scores for each of the selected conformations and pockets of *E. coli* cytochrome *bd*. (The notation for clusters is similar to Table 1.) The compound names are the ID numbers from the Life Chemicals molecule supplier.

The major point of using GaMD simulation to create a receptor ensemble for docking is that it can cover a large area of configuration space and dramatically improve the diversity of receptor structures.^54^ As shown in the Q-loop row of Table 1, the best docking score of different receptor structures varies considerably, about 29% from cluster 1 to cluster 4. Other pockets also show similar behavior. In addition, the top compounds are different for different clusters of one pocket (Tables 1 and 2), demonstrating the diversity of receptor structures. In sum, our GaMD simulations successfully enriched the receptor structural ensemble and greatly diversified the candidate molecules.

### 3. In vitro results

A collection of 102 compounds was selected for initial screening of inhibitory activity against *Mtb* cytochrome *bd* activity in inverted membrane vesicles energized with NADH as an electron donor for oxygen consumption. In these experiments, the *bcc:aa*_3_ oxidase complex was inhibited by TB47 (1 µM) so that all oxygen consumption was mediated by cytochrome *bd*. Of these compounds, 50 were obtained from molecular docking against the *E. coli bd* structure (Figure S6), 24 from the molecular docking of *Mtb* cytochrome *bd* with the disulfide bond (Figure S7), and 28 from the molecular docking against *Mtb* cytochrome *bd* without the disulfide bond (Figure S8). Compounds that inhibited OCR > 50% were selected for further testing. When Aurachin D (10 µM) was used as a positive control for inhibiting cytochrome *bd* activity, the OCR was inhibited approximately 90% (Figure S6).

From the *E. coli* set, three compounds inhibited the OCR > 50% (F1061-0249, F3259-0446, and F6617-3528) (Figure S6). From the two *Mtb* sets, only one compound from the molecular docking to cytochrome *bd* with the disulfide bond present set inhibited the OCR > 50% (F0777-1198) (Figure S7), while two compounds from molecular docking using cytochrome *bd* with a reduced disulfide bond present ((F2964-1180 and F3382-2259) were found to inhibit the OCR > 50% (Figure S8). These six compounds were then repeated in technical triplicate to define their activity more accurately (Figure 3 and Table 3). We also included the docked pose of the six hit compounds in Figure S4.

**Table 3.** Experimentally validated compounds with around and below 50% oxygen consumption rate. (Notation for docking structure: Mtb_disulfide stands for the structure from the trajectory of *M. tb* cytochrome *bd* with a disulfide bond (inactive conformation); *E. coli* stands for the structure from the trajectory of *E. coli* cytochrome *bd*; the rest of the notations are similar to Table 2. and Table 3.)

| Compounds | Docking scores | Docking structure | Rank in docking | OCR (%) |
| --- | --- | --- | --- | --- |
| F0777-1198 | -10.581 | Mtb_disulfide_mq9_4000 | 1 | 51.05% |
| F2964-1180 | -9.676 | Mtb_nodisulfide_oxy_190 | 2 | 50.63% |
| F3382-2259 | -10.068 | Mtb_nodisulfide_mq9_1412 | 2 | 30.41% |
| F1061-0249 | -12.952 | Ecoli_oxy_1170 | 6 | 31.82% |
| F3259-0446 | -12.903 | Ecoli_oxy_1170 | 7 | 32.27% |
| F6617-3528 | -13.072 | Ecoli_oxy_1846 | 4 | 18.99% |

**Figure 3.**
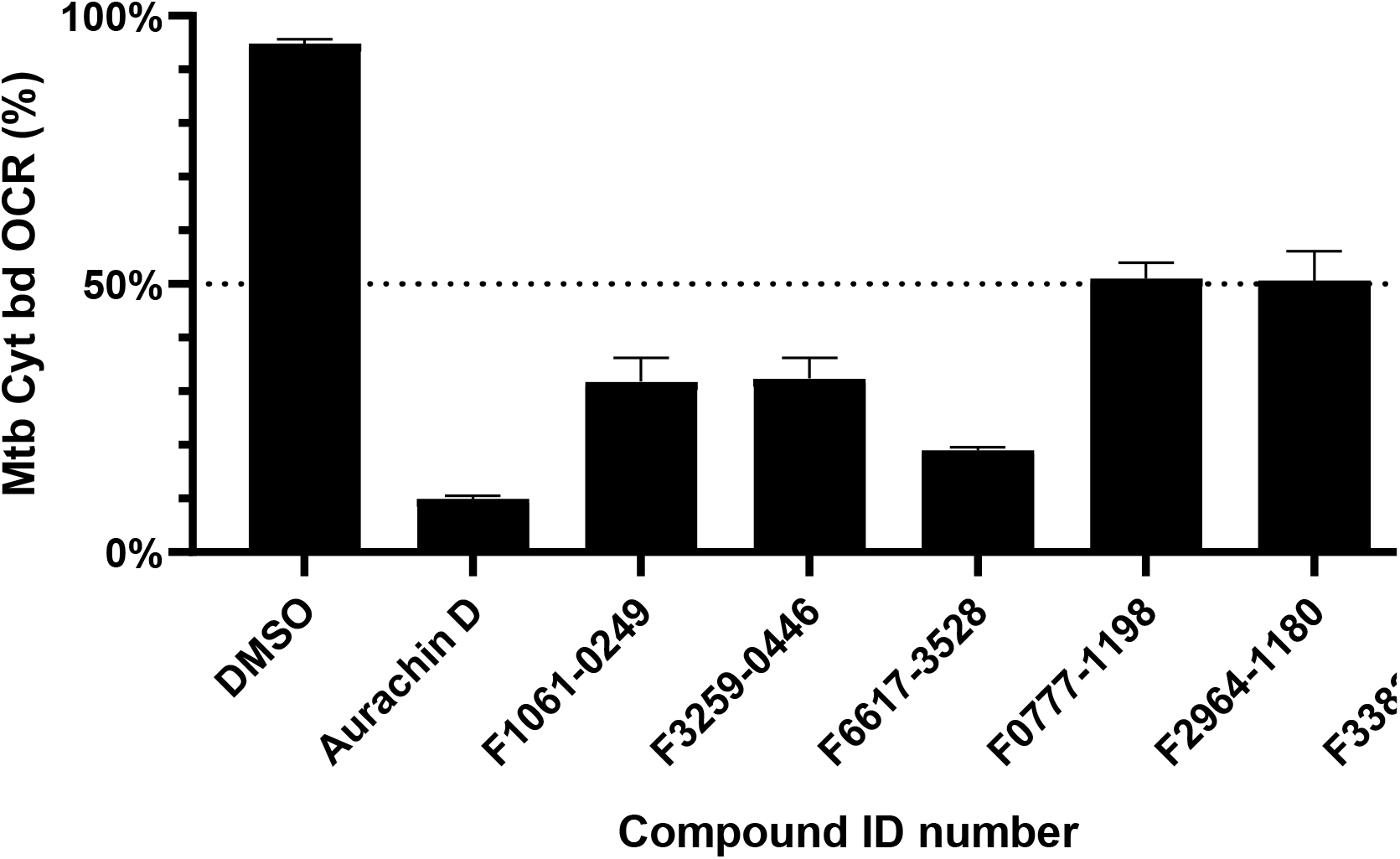
Remaining OCR of Mtb cytochrome bd in IMVs of M. smegmatis Δcyd pLHcyd after addition of compounds that passed initial screening at a concentration of 10 µM. IMVs were pre-treated with 1 µM TB47 for 2 minutes and energized with 500 µM NADH as the sole electron donor. DMSO vehicle control and Aurachin D (10 µM, > 90% inhibition) positive control are shown for comparison. Results are shown as the mean of technical triplicate with error bars showing standard deviation.

All compounds identified in the initial screen of the *E. coli* molecular docking were confirmed to be able to decrease the OCR to below 50% of initial levels, F1061-0249 at 31.82%, F3259-0446 at 32.37% and, F6617-3528 at 18.99% (Figure 4). From the *Mtb* molecular docking, only the compound F3382-2259 (from the molecular docking to the cytochrome *bd* with the reduced disulfide) decreased OCR below 50% of initial levels at 30.41% of base OCR. The compounds F0777-1198 and F2964-1180, which originated from the molecular docking with and without the disulfide bond present, respectively, were found to be unable to inhibit OCR to below 50%, with the mean activity after the addition of F0777-1198 being 51.05% and 50.63% after addition of F2964-1180. The compound F6617-3528 was the best inhibitor and the closest to the positive control Aurachin D, which inhibited OCR by 90%.

**Figure 4.**
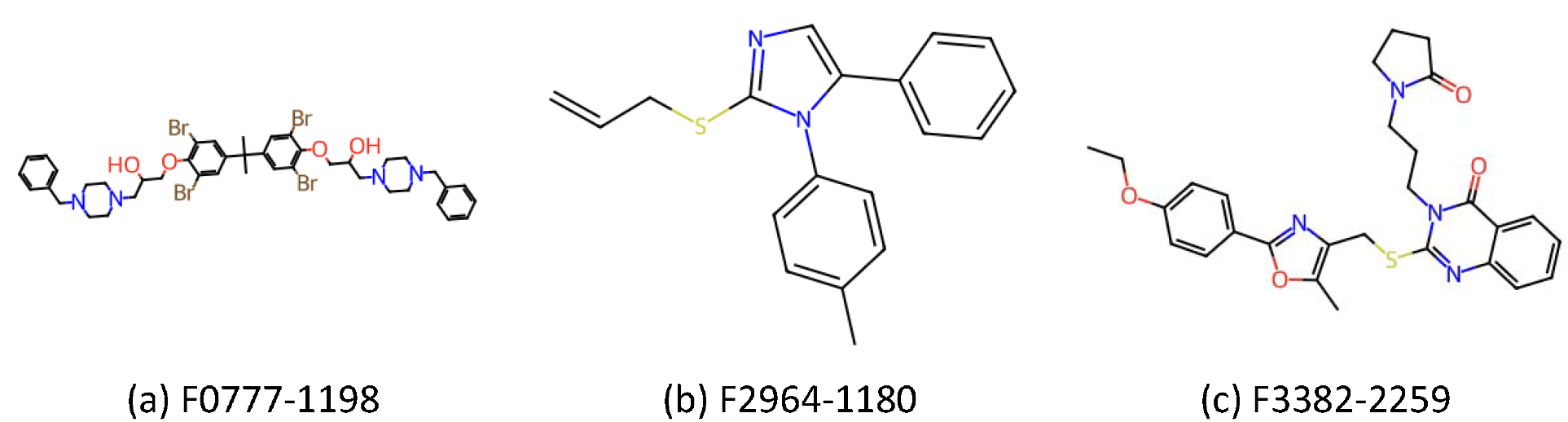
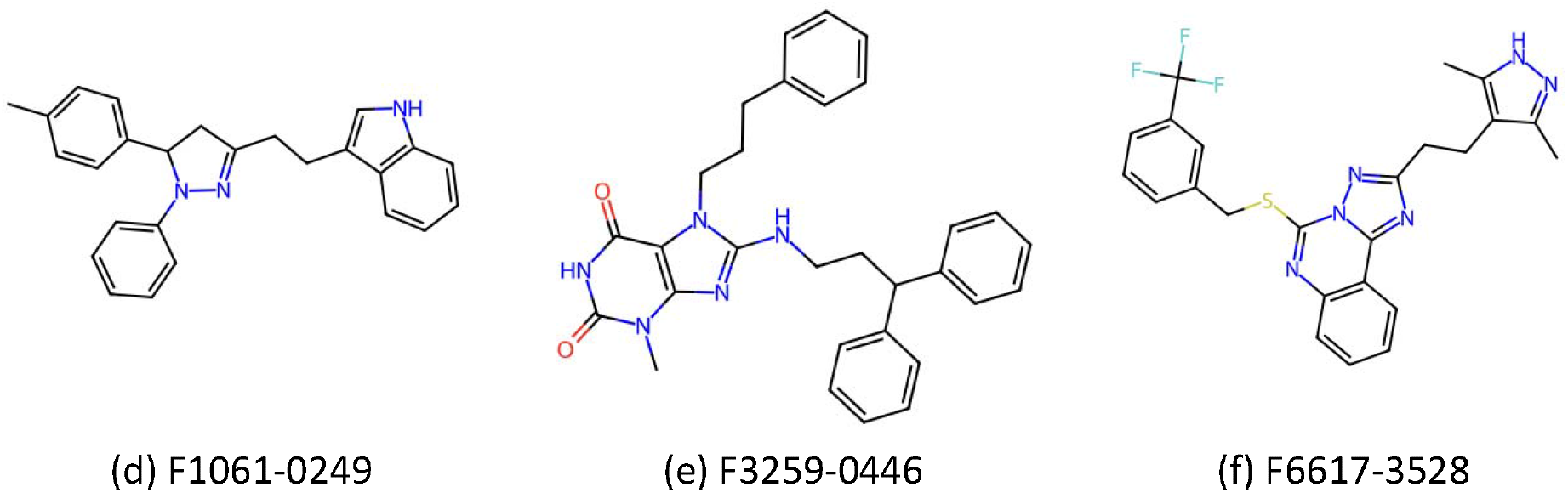
Structure of the experimentally validated compounds with around and below 50% oxygen consumption rate.

As seen from Table 3, four (F2964-1180, F1061-0249, F3259-0446, and F6617-3528) out of six hit compounds came from the same site around the head of the oxygen conducting channel. This observation can also be supported by the structural similarity of the four compounds. As shown in Figure 3, these four compounds have very similar shapes, resembling a triangle. Thus, all these four compounds may bind in the same pocket. The only exception in the hit compounds is F0777-1198. It was docked at the MQ9-binding site and shows a completely different structure than others (linear shape rather than triangular). It can be assumed that F0777-1198 represents a new binding mode, and its binding properties require further studies. The docking poses and protein-ligand interactions for all six hit compounds are shown in Figure S5.

## Conclusion

In this study, we used *in silico* techniques to generate and characterize potential drug-binding pockets and surfaces in *Mtb* and *E. coli* cytochrome *bd* before testing molecules as potential inhibitors against the *Mtb* cytochrome *bd*.

We employed GaMD simulations to create a variety of receptor conformations for ensemble docking. This method samples a larger free energy surface than traditional MD, enhancing the diversity of receptor structures. The variability in the best docking scores among different receptor structures, ranging about 29% from cluster 1 to cluster 4 for the Q-loop pocket, demonstrated the diversity of receptor structures.

To establish representative conformations from the GaMD trajectories, two clustering methods were employed: the DBScan clustering algorithm from the Amber CPPTRAJ program (for solvent-exposed binding surfaces, such as the Q-loop and QC-PL8 regions) and POVME 2.0 (for traditional pockets). Clustering results demonstrated distinct representative conformations of the pockets, and the utilization of different clustering methods increased the rigour of the analysis.

Between these two clustering algorithms, we selected 33 structures for ensemble docking, involving active and inactive conformations of *Mtb* cytochrome *bd* and *E. coli* cytochrome *bd*, employing the Schrödinger Maestro package. This enabled us to predict a diverse array of compounds for experimental tests.

A collection of 102 compounds underwent initial screening for inhibitory activity against *Mtb* cytochrome *bd*. Inverted membrane vesicles were employed, and the *bcc:aa*_3_ oxidase complex was inhibited by TB47, ensuring that all oxygen consumption measurements were mediated by cytochrome *bd*. Compounds inhibiting OCR by > 50% were selected for further testing.

The experiments showed that three compounds screened *in silico* against *E. coli* (F1061-0249, F3259-0446, and F6617-3528) significantly reduced OCR. From the molecules screened *in silico* against *Mtb*, only one compound (F0777-1198) from the oxidized disulfide bond set (active conformation) lowered OCR below 50%, while two compounds (F2964-1180 and F3382-2259) from the reduced disulfide bond set (inactive conformation) exhibited around 50% inhibitory effects. These findings were subjected to triplicate testing, revealing F6617-3528 as the most effective inhibitor, closely resembling the positive control Aurachin D. These six structures do not resemble the canonical inhibitor Aurachin D or other known inhibitors, offering promising avenues for new drug development towards cytochrome *bd*.

When analyzing the hit compounds, four (F2964-1180, F1061-0249, F3259-0446, and F6617-3528) of six compounds shared a similar structural site around the head of the oxygen-conducting channel. This observation was supported by their structural resemblance, a triangular shape, suggesting a common binding pose. An interesting exception was F0777-1198, which was docked at the MQ9 binding site with a distinct linear structure and may represent a potential new binding mode. Compound F3382-2259 also docked to the MQ9 binding site. However, these binding poses were not evaluated experimentally, so the exact binding site of these inhibitors may differ from our expectations.

In summary, this study successfully characterized potential drug-binding pockets of Mtb cytochrome *bd* oxidase, with GaMD simulations to diversify receptor structures for docking. The predicted new inhibitors were tested through inhibitory activity assays. The findings contribute valuable insights into potential drug candidates for targeting cytochrome *bd* oxidases in *Mtb* and *E. coli*. The identified hit compounds, especially F6617-3528, hold promise for further exploration and development as potential drugs against these bacteria.

## Acknowledgement

C.S. was supported by a National Science Foundation Graduate Research Fellowship (DGE-1650112). S.A. acknowledges support from the Department of Chemical Engineering at the University of California, Davis. This work used Expanse at San Diego Supercomputer Center (SDSC) through allocation BIO210051 to S.A. from the Extreme Science and Engineering Discovery Environment (XSEDE), which was supported by National Science Foundation grant number #1548562.^55^ KLK and GMC acknowledge support from the Health Research Council, Grant 23-396 the Marsden Fund UOO1603, and the RSNZ Catalyst Fund UOO1702.

## References

(1) Borisov, V. B.; Siletsky, S. A.; Paiardini, A.; Hoogewijs, D.; Forte, E.; Giuffre, A.; Poole, R. K. Bacterial oxidases of the cytochrome bd family: Redox enzymes of unique structure, function, and utility as drug targets. Antioxidants & redox signaling 2021, 34 (16), 1280–1318.

(2) Cook, G. M.; Poole, R. K. A bacterial oxidase like no other? Science 2016, 352 (6285), 518–519.

(3) Friedrich, T.; Wohlwend, D.; Borisov, V. B. Recent advances in structural studies of cytochrome bd and its potential application as a drug target. International Journal of Molecular Sciences 2022, 23 (6), 3166.

(4) Yaoi, H.; Tamiya, H. On the respiratory pigment, cytochrome, in bacteria. Proceedings of the Imperial Academy 1928, 4 (7), 436–439.

(5) Safarian, S.; Rajendran, C.; Müller, H.; Preu, J.; Langer, J. D.; Ovchinnikov, S.; Hirose, T.; Kusumoto, T.; Sakamoto, J.; Michel, H. Structure of a bd oxidase indicates similar mechanisms for membrane-integrated oxygen reductases. Science 2016, 352 (6285), 583–586.

(6) Safarian, S.; Hahn, A.; Mills, D.; Radloff, M.; Eisinger, M. L.; Nikolaev, A.; Meier-Credo, J.; Melin, F.; Miyoshi, H.; Gennis, R. Active site rearrangement and structural divergence in prokaryotic respiratory oxidases. Science 2019, 366 (6461), 100–104.

(7) Theßeling, A.; Rasmussen, T.; Burschel, S.; Wohlwend, D.; Kägi, J.; Müller, R.; Böttcher, B.; Friedrich, T. Homologous bd oxidases share the same architecture but differ in mechanism. Nature communications 2019, 10 (1), 5138.

(8) Grauel, A.; Kägi, J.; Rasmussen, T.; Makarchuk, I.; Oppermann, S.; Moumbock, A. F.; Wohlwend, D.; Müller, R.; Melin, F.; Günther, S. Structure of Escherichia coli cytochrome bd-II type oxidase with bound aurachin D. Nature Communications 2021, 12 (1), 6498.

(9) Grund, T. N.; Radloff, M.; Wu, D.; Goojani, H. G.; Witte, L. F.; Jösting, W.; Buschmann, S.; Müller, H.; Elamri, I.; Welsch, S. Mechanistic and structural diversity between cytochrome bd isoforms of Escherichia coli. Proceedings of the National Academy of Sciences 2021, 118 (50), e2114013118.

(10) Safarian, S.; Opel-Reading, H. K.; Wu, D.; Mehdipour, A. R.; Hards, K.; Harold, L. K.; Radloff, M.; Stewart, I.; Welsch, S.; Hummer, G. The cryo-EM structure of the bd oxidase from M. tuberculosis reveals a unique structural framework and enables rational drug design to combat TB. Nature Communications 2021, 12 (1), 5236.

(11) Wang, W.; Gao, Y.; Tang, Y.; Zhou, X.; Lai, Y.; Zhou, S.; Zhang, Y.; Yang, X.; Liu, F.; Guddat, L. W. Cryo-EM structure of mycobacterial cytochrome bd reveals two oxygen access channels. Nature Communications 2021, 12 (1), 4621.

(12) Meunier, B.; Madgwick, S. A.; Reil, E.; Oettmeier, W.; Rich, P. R. New inhibitors of the quinol oxidation sites of bacterial cytochromes bo and bd. Biochemistry 1995, 34 (3), 1076–1083.

(13) Lu, P.; Heineke, M. H.; Koul, A.; Andries, K.; Cook, G. M.; Lill, H.; van Spanning, R.; Bald, D. The cytochrome bd-type quinol oxidase is important for survival of Mycobacterium smegmatis under peroxide and antibiotic-induced stress. Scientific reports 2015, 5 (1), 10333.

(14) Radloff, M.; Elamri, I.; Grund, T. N.; Witte, L. F.; Hohmann, K. F.; Nakagaki, S.; Goojani, H. G.; Nasiri, H.; Miyoshi, H.; Bald, D. Short-chain aurachin D derivatives are selective inhibitors of E. coli cytochrome bd-I and bd-II oxidases. Scientific Reports 2021, 11 (1), 23852.

(15) Miyoshi, H.; Takegami, K.; Sakamoto, K.; Mogi, T.; Iwamura, H. Characterization of the ubiquinol oxidation sites in cytochromes bo and bd from Escherichia coli using aurachin C analogues. The journal of biochemistry 1999, 125 (1), 138–142.

(16) Jeffreys, L. N.; Ardrey, A.; Hafiz, T. A.; Dyer, L.-A.; Warman, A. J.; Mosallam, N.; Nixon, G. L.; Fisher, N. E.; Hong, W. D.; Leung, S. C. Identification of 2-Aryl-Quinolone Inhibitors of Cytochrome bd and Chemical Validation of Combination Strategies for Respiratory Inhibitors against Mycobacterium tuberculosis. ACS Infectious Diseases 2023, 9 (2), 221–238.

(17) Zhou, Y.; Shao, M.; Wang, W.; Cheung, C.-Y.; Wu, Y.; Yu, H.; Hu, X.; Cook, G. M.; Gong, H.; Lu, X. Discovery of 1-hydroxy-2-methylquinolin-4 (1H)-one derivatives as new cytochrome bd oxidase inhibitors for tuberculosis therapy. European Journal of Medicinal Chemistry 2023, 245, 114896.

(18) Kumar, A.; Kumari, N.; Bhattacherjee, S.; Venugopal, U.; Parwez, S.; Siddiqi, M. I.; Krishnan, M. Y.; Panda, G. Design, synthesis and biological evaluation of (Quinazoline 4-yloxy) acetamide and (4-oxoquinazoline-3 (4H)-yl) acetamide derivatives as inhibitors of Mycobacterium tuberculosis bd oxidase. European Journal of Medicinal Chemistry 2022, 242, 114639.

(19) Lawer, A.; Tyler, C.; Hards, K.; Keighley, L. M.; Cheung, C.-Y.; Kierek, F.; Su, S.; Matikonda, S. S.; McInnes, T.; Tyndall, J. D. Synthesis and biological evaluation of aurachin D analogues as inhibitors of Mycobacterium tuberculosis cytochrome bd oxidase. ACS Medicinal Chemistry Letters 2022, 13 (10), 1663–1669.

(20) Makarchuk, I.; Nikolaev, A.; Thesseling, A.; Dejon, L.; Lamberty, D.; Stief, L.; Speicher, A.; Friedrich, T.; Hellwig, P.; Nasiri, H. Identification and optimization of quinolone-based inhibitors against cytochrome bd oxidase using an electrochemical assay. Electrochimica Acta 2021, 381, 138293.

(21) Harikishore, A.; Chong, S. S. M.; Ragunathan, P.; Bates, R. W.; Grüber, G. Targeting the menaquinol binding loop of mycobacterial cytochrome bd oxidase. Molecular diversity 2021, 25, 517–524.

(22) Hopfner, S. M.; Lee, B. S.; Kalia, N. P.; Miller, M. J.; Pethe, K.; Moraski, G. C. Structure guided generation of thieno [3, 2-d] pyrimidin-4-amine Mycobacterium tuberculosis bd oxidase inhibitors. RSC Medicinal Chemistry 2021, 12 (1), 73–77.

(23) Lee, B. S.; Hards, K.; Engelhart, C. A.; Hasenoehrl, E. J.; Kalia, N. P.; Mackenzie, J. S.; Sviriaeva, E.; Chong, S. M. S.; Manimekalai, M. S. S.; Koh, V. H. Dual inhibition of the terminal oxidases eradicates antibiotic-tolerant Mycobacterium tuberculosis. EMBO Molecular Medicine 2021, 13 (1), e13207.

(24) Hards, K.; Cheung, C.-Y.; Waller, N.; Adolph, C.; Keighley, L.; Tee, Z. S.; Harold, L. K.; Menorca, A.; Bujaroski, R. S.; Buckley, B. J. An amiloride derivative is active against the F1Fo-ATP synthase and cytochrome bd oxidase of Mycobacterium tuberculosis. Communications Biology 2022, 5 (1), 166.

(25) Saha, P.; Das, S.; Indurthi, H. K.; Kumar, R.; Roy, A.; Kalia, N. P.; Sharma, D. K. Cytochrome bd oxidase: an emerging anti-tubercular drug target. RSC Medicinal Chemistry 2024. DOI: 10.1039/D3MD00587A (acccessed February 23, 2024).

(26) Harikishore, A.; Mathiyazakan, V.; Pethe, K.; Grüber, G. Novel targets and inhibitors of the Mycobacterium tuberculosis cytochrome bd oxidase to foster anti-tuberculosis drug discovery. Expert Opinion on Drug DiscSovery 2023, 1–11.

(27) Miao, Y.; Feher, V. A.; McCammon, J. A. Gaussian accelerated molecular dynamics: Unconstrained enhanced sampling and free energy calculation. Journal of chemical theory and computation 2015, 11 (8), 3584–3595.

(28) Miao, Y.; McCammon, J. A. Gaussian accelerated molecular dynamics: Theory, implementation, and applications. In Annual reports in computational chemistry, Vol. 13; Elsevier, 2017; pp 231–278.

(29) Wang, J.; Arantes, P. R.; Bhattarai, A.; Hsu, R. V.; Pawnikar, S.; Huang, Y. m. M.; Palermo, G.; Miao, Y. Gaussian accelerated molecular dynamics: Principles and applications. Wiley Interdisciplinary Reviews: Computational Molecular Science 2021, 11 (5), e1521.

(30) Ahn, S.-H.; Seitz, C.; Cruzeiro, V. W. D.; McCammon, J. A.; Götz, A. W. Data for molecular dynamics simulations of Escherichia coli cytochrome bd oxidase with the Amber force field. Data in Brief 2021, 38, 107401.

(31) Schrödinger Release 2023-3: Maestro. 2023; Schrödinger, LLC, New York, NY.

(32) Olsson, M. H.; Søndergaard, C. R.; Rostkowski, M.; Jensen, J. H. PROPKA3: consistent treatment of internal and surface residues in empirical p K a predictions. Journal of chemical theory and computation 2011, 7 (2), 525–537.

(33) Jumper, J.; Evans, R.; Pritzel, A.; Green, T.; Figurnov, M.; Ronneberger, O.; Tunyasuvunakool, K.; Bates, R.; Žídek, A.; Potapenko, A. Highly accurate protein structure prediction with AlphaFold. Nature 2021, 596 (7873), 583–589.

(34) Case, D. A.; Aktulga, H. M.; Belfon, K.; Ben-Shalom, I.; Brozell, S. R.; Cerutti, D. S.; Cheatham III, T. E.; Cruzeiro, V. W. D.; Darden, T. A.; Duke, R. E. Amber 2021; University of California, San Francisco, 2021.

(35) Brooks, B. R.; Brooks III, C. L.; Mackerell Jr, A. D.; Nilsson, L.; Petrella, R. J.; Roux, B.; Won, Y.; Archontis, G.; Bartels, C.; Boresch, S. CHARMM: the biomolecular simulation program. Journal of computational chemistry 2009, 30 (10), 1545–1614.

(36) Jo, S.; Kim, T.; Iyer, V. G.; Im, W. CHARMM-GUI: a web-based graphical user interface for CHARMM. Journal of computational chemistry 2008, 29 (11), 1859–1865.

(37) Morein, S.; Andersson, A.-S.; Rilfors, L.; Lindblom, G. Wild-type Escherichia coli Cells Regulate the Membrane Lipid Composition in a “Window” between Gel and Non-lamellar Structures (*). Journal of Biological Chemistry 1996, 271 (12), 6801–6809.

(38) Kumar, G.; Kalra, V. K.; Brodie, A. F. Asymmetric distribution of phospholipids in membranes from Mycobacterium phlei. Archives of Biochemistry and Biophysics 1979, 198 (1), 22–30.

(39) Daffé, M.; Draper, P. The envelope layers of mycobacteria with reference to their pathogenicity. Advances in microbial physiology 1997, 39, 131–203.

(40) Jackson, M.; Crick, D. C.; Brennan, P. J. Phosphatidylinositol is an essential phospholipid of mycobacteria. Journal of Biological Chemistry 2000, 275 (39), 30092–30099.

(41) Daffé, M.; Marrakchi, H. Unraveling the structure of the mycobacterial envelope. Microbiology spectrum 2019, 7 (4), 7.4.1.

(42) Jorgensen, W. L.; Chandrasekhar, J.; Madura, J. D.; Impey, R. W.; Klein, M. L. Comparison of simple potential functions for simulating liquid water. The Journal of chemical physics 1983, 79 (2), 926–935.

(43) Beglov, D.; Roux, B. Finite representation of an infinite bulk system: solvent boundary potential for computer simulations. The Journal of chemical physics 1994, 100 (12), 9050–9063.

(44) Maier, J. A.; Martinez, C.; Kasavajhala, K.; Wickstrom, L.; Hauser, K. E.; Simmerling, C. ff14SB: improving the accuracy of protein side chain and backbone parameters from ff99SB. Journal of chemical theory and computation 2015, 11 (8), 3696–3713.

(45) Gould, I.; Skjevik, A.; Dickson, C.; Madej, B.; Walker, R. Lipid17: A comprehensive AMBER force field for the simulation of zwitterionic and anionic lipids. Manuscript in preparation 2018.

(46) Wang, J.; Wolf, R. M.; Caldwell, J. W.; Kollman, P. A.; Case, D. A. Development and testing of a general amber force field. Journal of computational chemistry 2004, 25 (9), 1157–1174.

(47) Schrödinger, LLC.

(48) Durrant, J. D.; Votapka, L.; Sørensen, J.; Amaro, R. E. POVME 2.0: an enhanced tool for determining pocket shape and volume characteristics. Journal of chemical theory and computation 2014, 10 (11), 5047–5056.

(49) Schrödinger Release 2022-1: Protein Preparation Wizard. 2022; Schrödinger, LLC, New York, NY.

(50) Schrödinger Release 2022-1: Epik. 2022; Schrödinger, LLC, New York, NY.

(51) Schrödinger Release 2022-1: Prime. 2022; Schrödinger, LLC, New York, NY.

(52) Schrödinger Release 2022-1: Glide. 2022; Schrödinger, LLC, New York, NY.

(53) Lu, X.; Williams, Z.; Hards, K.; Tang, J.; Cheung, C.-Y.; Aung, H. L.; Wang, B.; Liu, Z.; Hu, X.; Lenaerts, A. Pyrazolo [1, 5-a] pyridine inhibitor of the respiratory cytochrome bcc complex for the treatment of drug-resistant tuberculosis. ACS infectious diseases 2018, 5 (2), 239–249.

(54) Miao, Y.; Goldfeld, D. A.; Moo, E. V.; Sexton, P. M.; Christopoulos, A.; McCammon, J. A.; Valant, C. Accelerated structure-based design of chemically diverse allosteric modulators of a muscarinic G protein-coupled receptor. Proceedings of the National Academy of Sciences 2016, 113 (38), E5675–E5684.

(55) Towns, J.; Cockerill, T.; Dahan, M.; Foster, I.; Gaither, K.; Grimshaw, A.; Hazlewood, V.; Lathrop, S.; Lifka, D.; Peterson, G. D.; et al. XSEDE: Accelerating Scientific Discovery. Comput. Sci. Eng. September-October 2014 16, 62–74. 10.1109/MCSE.2014.80 (acccessed June 17, 2020).

