## Supplementary Materials for "Targeting Tuberculosis: Novel Scaffolds for Inhibiting Cytochrome bd Oxidase"

∥*These authors have contributed equally.*

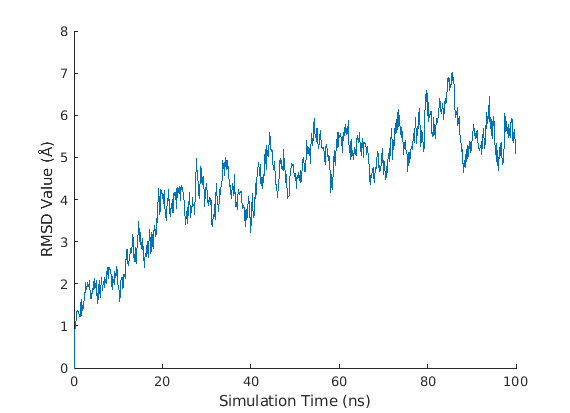

(a)

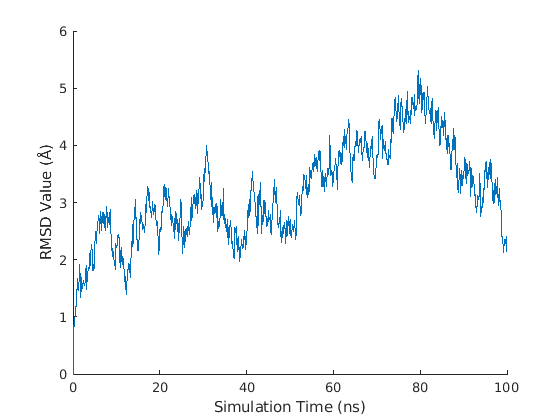

(b)

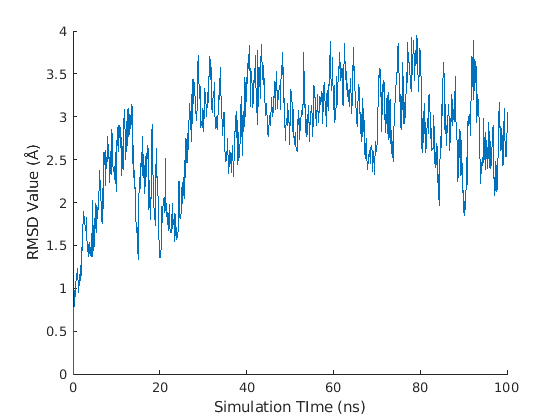

(c)

Figure S1. Root-mean-square deviation (RMSD) plot of different cytochrome bd system relative to its initial conformation during a 100 ns conventional MD simulation. (a) E. coli cytochrome bd system (b) Mtb cytochrome bd system with the disulfied bond (c) Mtb cytochrome bd system without the disulfied bond

| 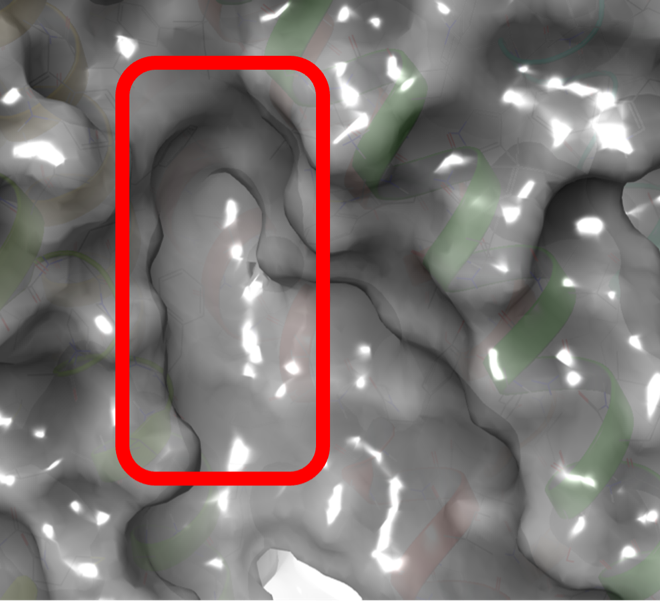 | 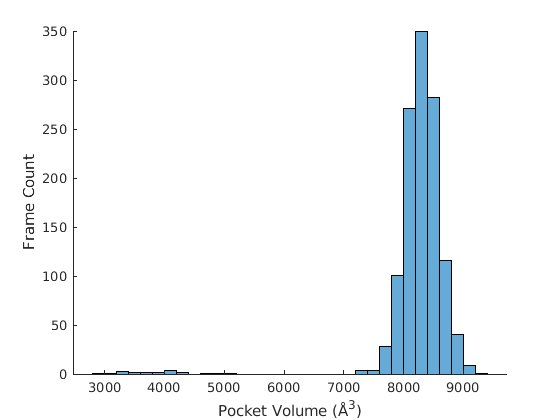 |
| --- | --- |
| (a) | (b) |
| 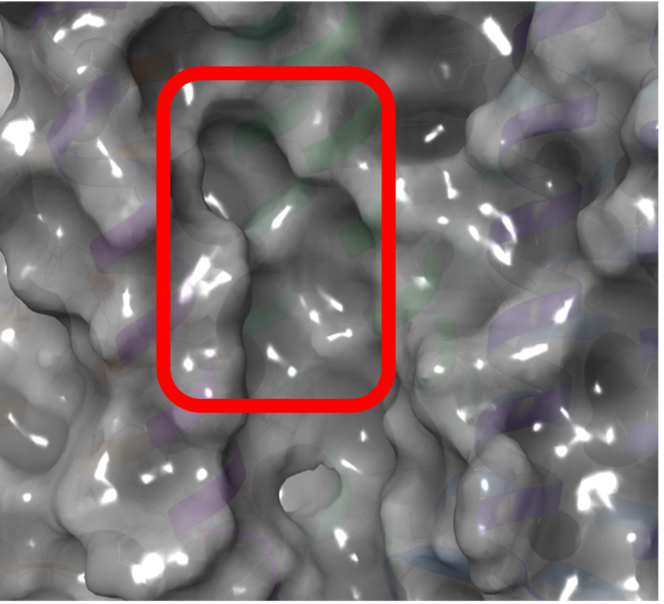 | 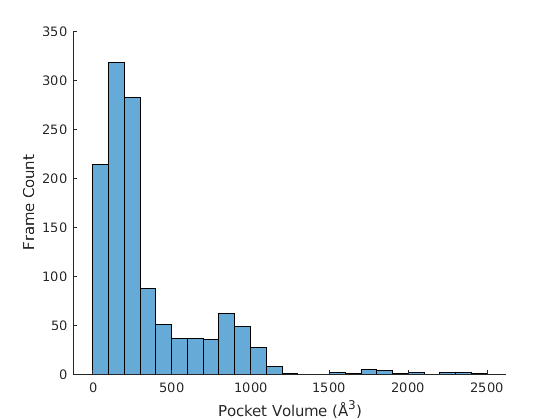 |
| (c) | (d) |
| 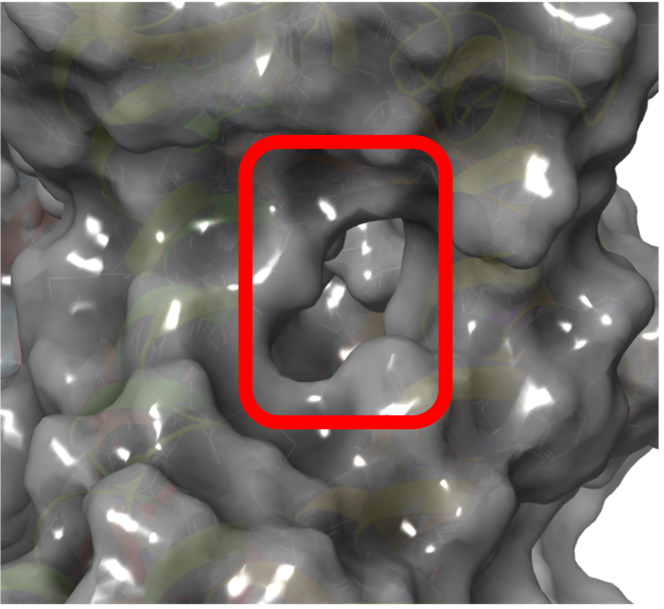 | 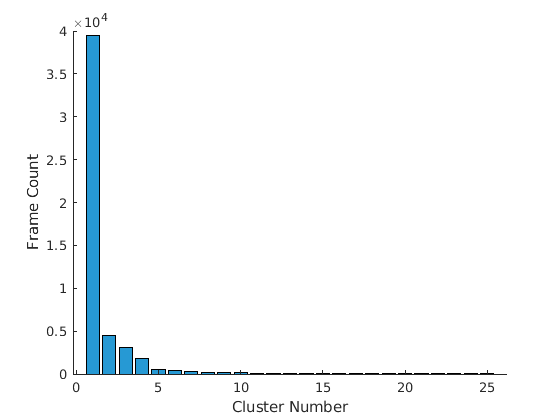 |
| (e) | (f) |
| 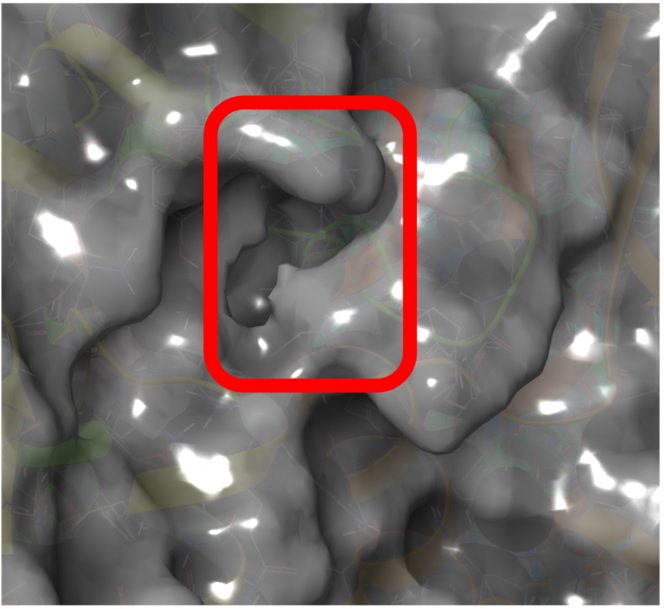 | |
| (g) | |

Figure S2. Representative pocket conformations and the population distribution for GaMD trajectories of *Mtb* cytochrome *bd* with the disulfide bond (inactive conformation). (a) MQ9 binding site and (b) pocket volume distribution of it; (c) oxygen conducting channel and (d) pocket volume distribution of it; (e) the Q-loop region and (f) cluster population distribution of it; (g) the region between QC and PL8 (There is no cluster population distribution because there is only one representative structure).

| 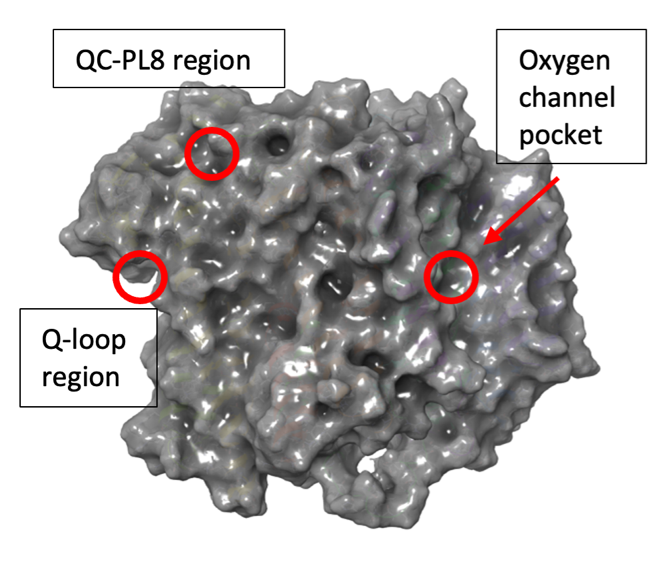 | 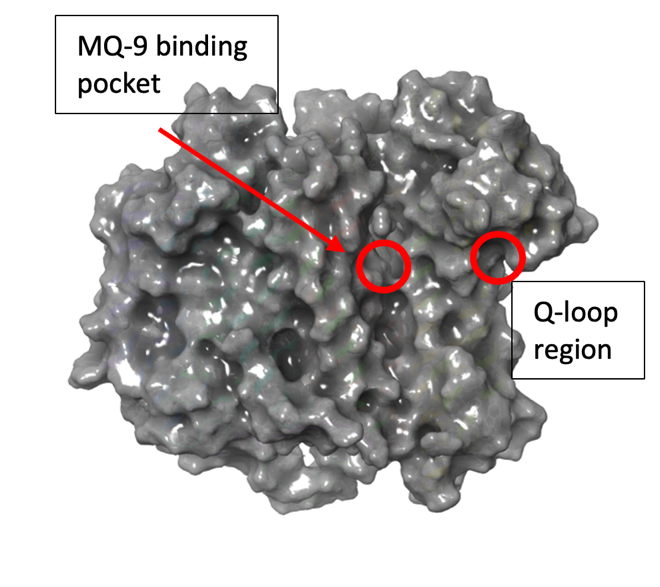 |
| --- | --- |
| (a) | (b) |
| 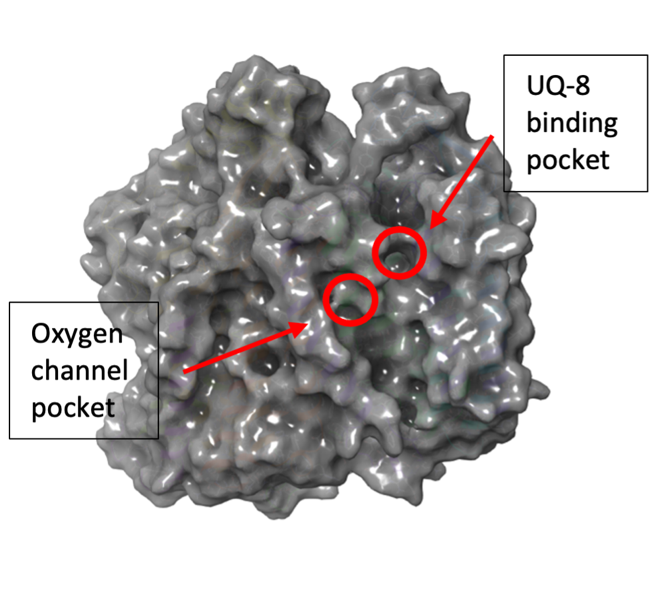 | 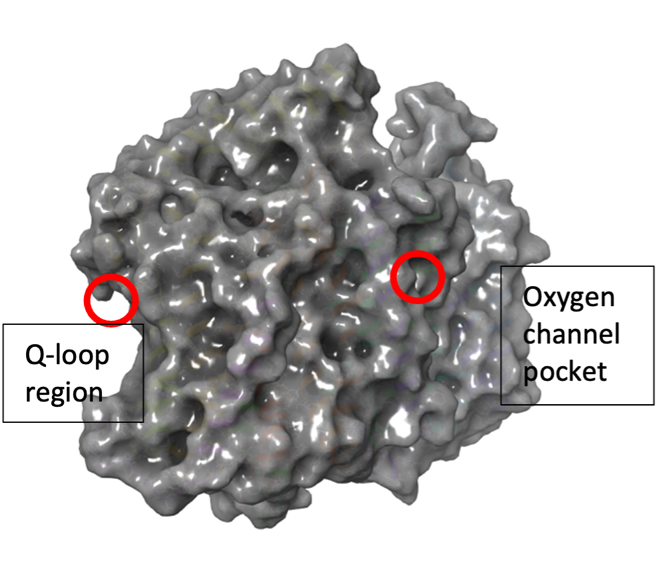 |
| (c) | (d) |

Figure S3. Location of the chosen pockets. (a) and (b) are for the *Mtb* cytochrome *bd* oxidase, and (c) and (d) are for the *E. coli* cytochrome *bd* oxidase.

| 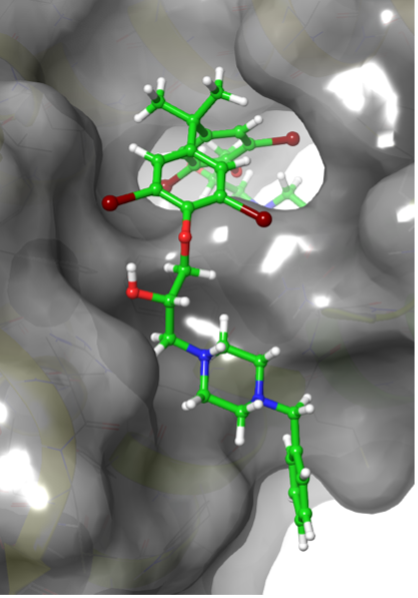 | 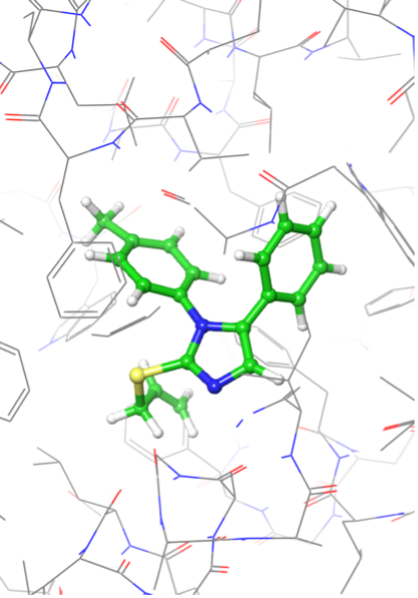 | 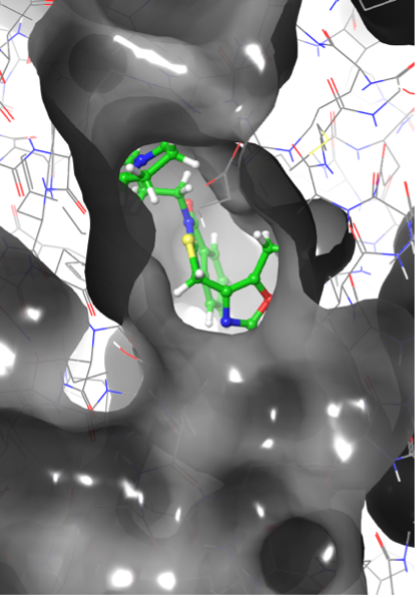 |
| --- | --- | --- |
| (a) F0777-1198 | (b) F2964-1180 | (c) F3382-2259 |
| 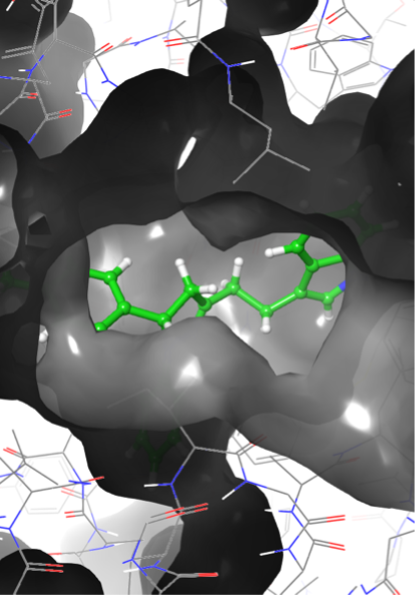 | 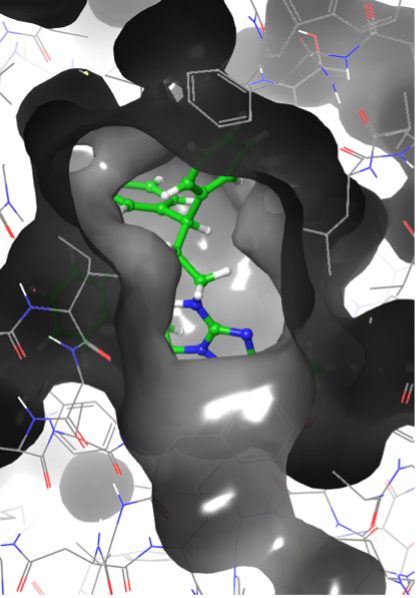 | 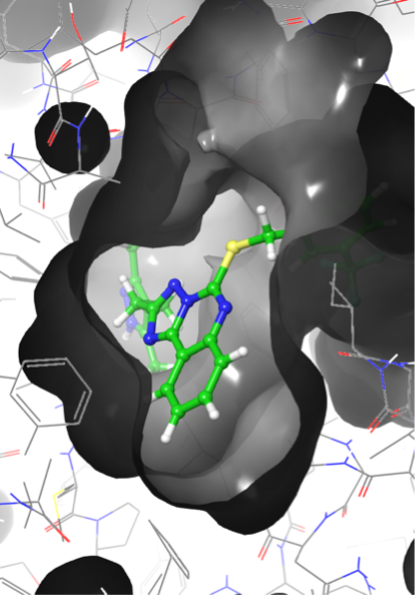 |
| (d) F1061-0249 | (e) F3259-0446 | (f) F6617-3528 |

Figure S4. Docking poses of the hit compounds. (a) compound F0777-1198 docked to the MQ9 binding site in the active Mtb conformation. (b) compound F2964-1180 docked to the oxygen channel in the inactive Mtb conformation. (c) compound F3382-2259 docked to the MQ9 binding site in the inactive Mtb conformation. (d) compound F1061-0249 docked to the oxygen channel in E. coli. (e) compound F3259-0446 docked to the oxygen channel in E. coli. (f) compound F6617-3528 docked to the oxygen channel in E. coli.

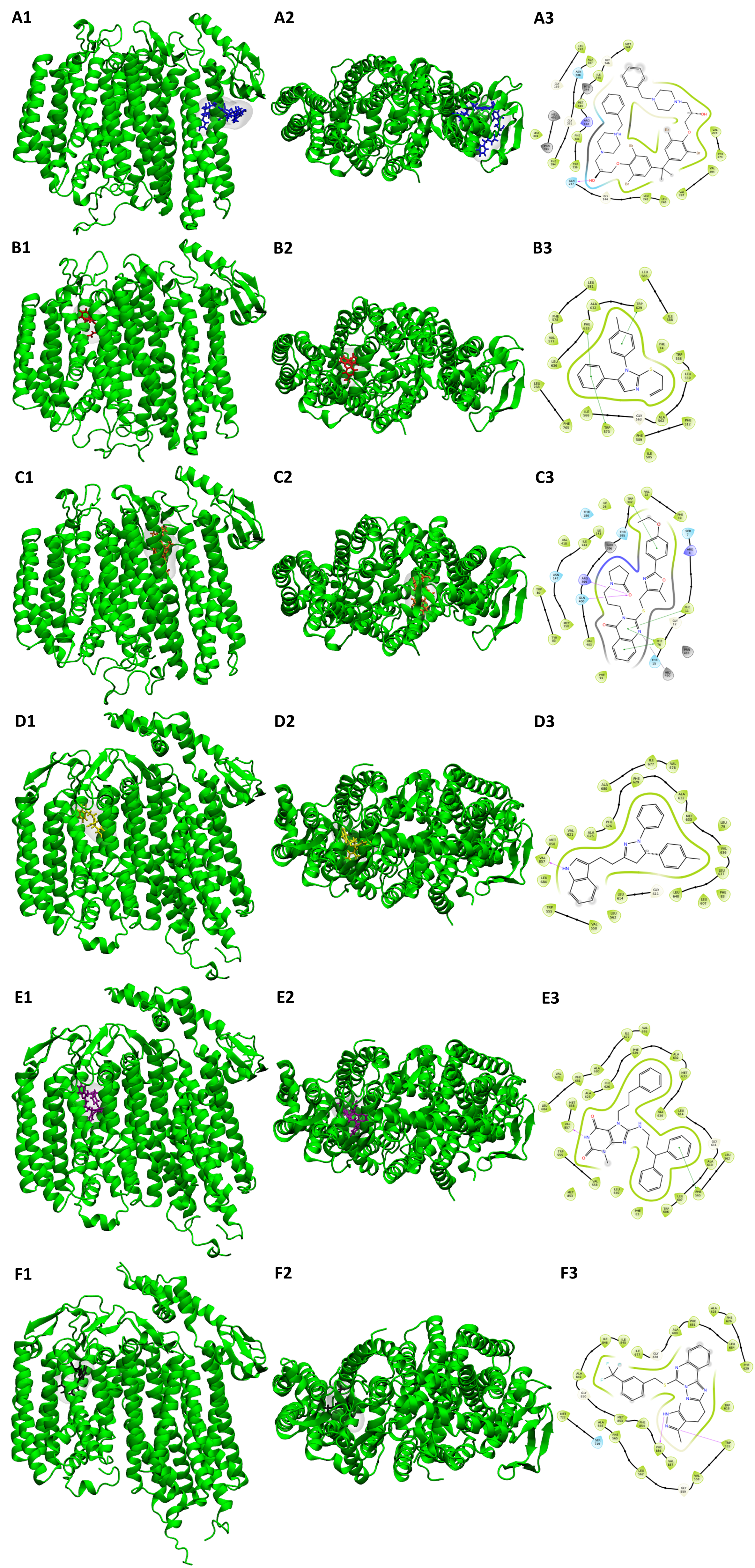

Figure S5. Docking poses and ligand interactions. To better visualize the locations and protein-ligand interactions of our hits, we show (1) A side view of the docked ligand in the protein (2) A top view of the docked ligand in the protein (3) the protein-ligand interactions for each of our hit ligands: (A) F0777-1198 in blue (B) F2964-1180 in red (C) F3382-2259 in orange (D) F1061-0249 in yellow (E) F3259-0446 in purple (F) F6617-3528 in black. Cytochrome bd is in green.

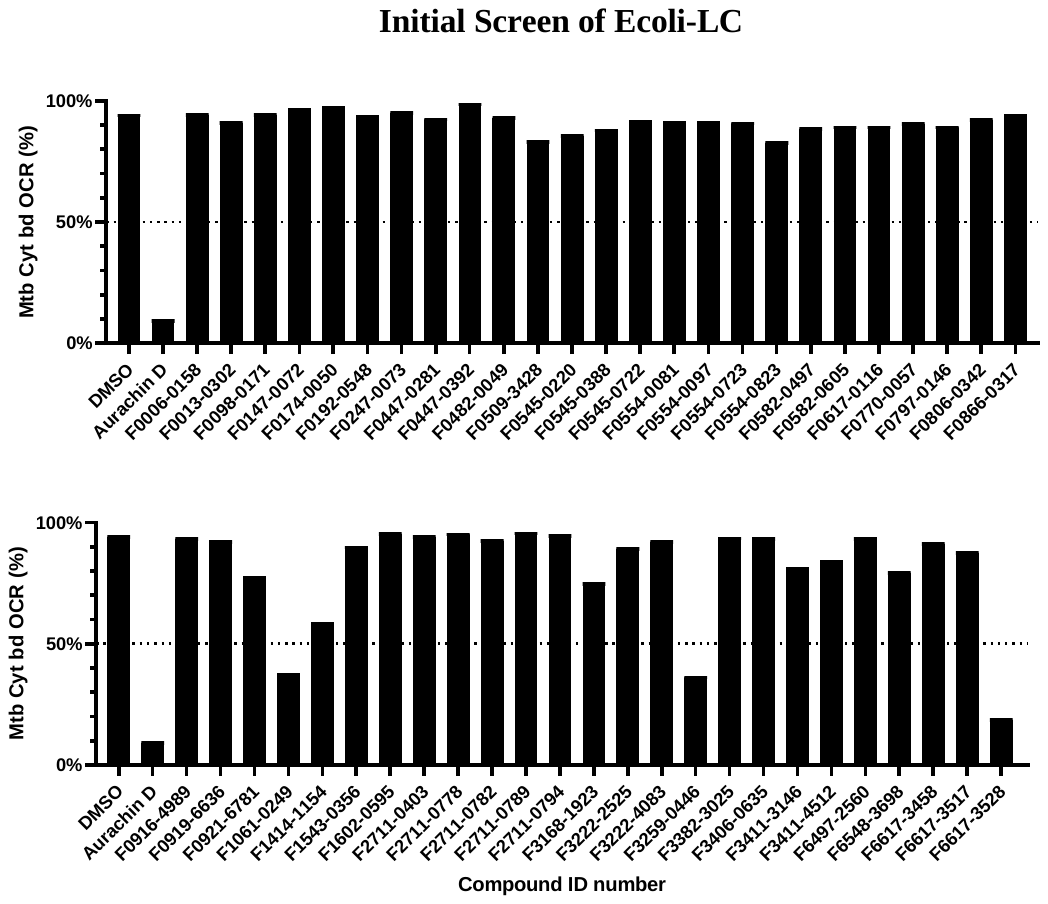

Figure S6. Single replicate screen showing remaining OCR of Mtb cytochrome bd oxidase in IMVs of M. smegmatis Δcyd pLHcyd after addition of compounds from E. coli list at a concentration of 10 µM. IMVs were pre-treated with 1 µM TB47 and energised with 500 µM NADH as the sole electron donor. DMSO vehicle control and Aurachin D (10 µM) positive control are shown for comparison.

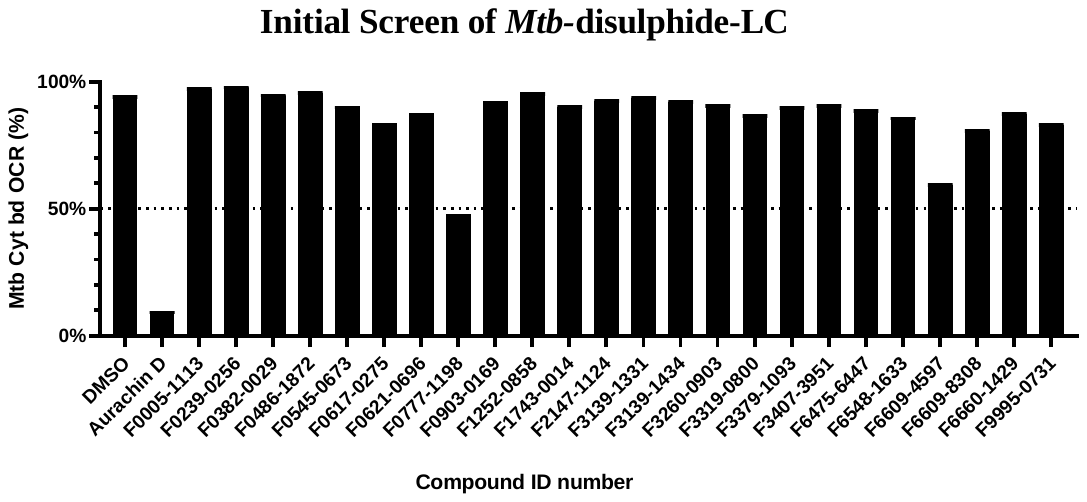

Figure S7. Single replicate screen showing remaining OCR of Mtb cytochrome bd oxidase in IMVs of M. smegmatis Δcyd pLHcyd after addition of compounds from Mtb-disulfide list at a concentration of 10 µM. IMVs were pre-treated with 1 µM TB47 and energised with 500 µM NADH as the sole electron donor. DMSO vehicle control and Aurachin D (10 µM) positive control are shown for comparison.

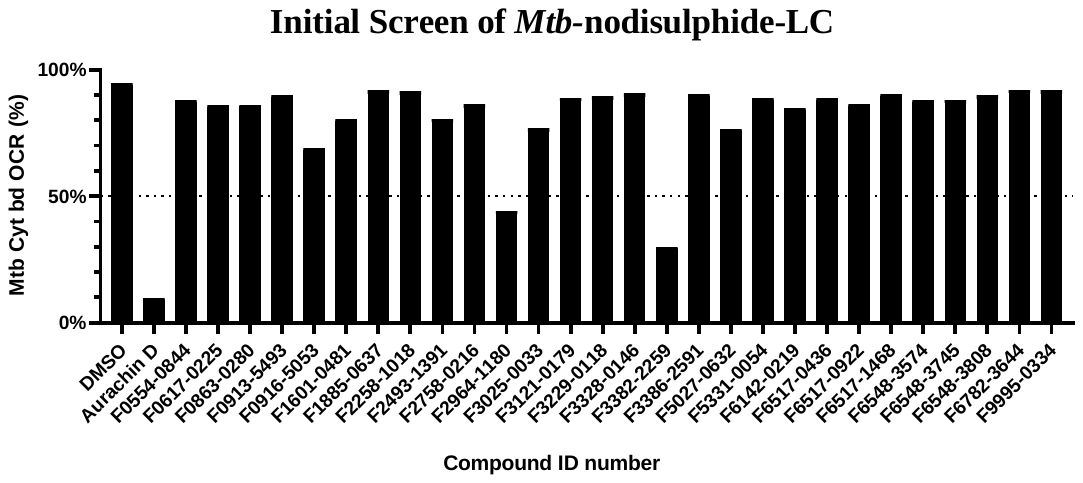

Figure S8. Single replicate screen showing remaining OCR of Mtb cytochrome bd oxidase in IMVs of M. smegmatis Δcyd pLHcyd after addition of compounds from Mtb-no disulfide list at a concentration of 10 µM. IMVs were pre-treated with 1 µM TB47 and energised with 500 µM NADH as the sole electron donor. DMSO vehicle control and Aurachin D (10 µM) positive control are shown for comparison.
